# The impact of di-2-ethylhexyl phthalate on sperm fertility

**DOI:** 10.1101/2020.04.09.034835

**Authors:** Liliya Gabelev Khasin, John Della Rosa, Natalie Petersen, Jacob Moeller, Lance J. Kriegsfeld, Polina V. Lishko

## Abstract

A growing number of studies point to reduced fertility upon chronic exposure to endocrine-disrupting chemicals (EDCs) such as phthalates and plasticizers. These toxins are ubiquitous and are often found in food and beverage containers, medical devices, as well as in common household and personal care items. Animal studies with EDCs, such as phthalates and bisphenol A have shown a dose-dependent decrease in fertility and embryo toxicity upon chronic exposure. However, limited research has been conducted on the acute effects of these EDCs on male fertility. Here we used a murine model to test the acute effects of four ubiquitous environmental toxins: bisphenol A (BPA), di-2-ethylhexyl phthalate (DEHP), diethyl phthalate (DEP), and dimethyl phthalate (DMP) on sperm fertilizing ability and pre-implantation embryo development. The most potent of these toxins, di-2-ethylhexyl phthalate (DEHP), was further evaluated for its effect on sperm ion channel activity, capacitation status, acrosome reaction and generation of reactive oxygen species (ROS). DEHP demonstrated a profound hazardous effect on sperm fertility by producing an altered capacitation profile, impairing the acrosome reaction, and, interestingly, also increasing ROS production. These results indicate that in addition to its known chronic impact on reproductive potential, DEHP also imposes acute and profound damage to spermatozoa, and thus, represents a significant risk to male fertility.

## Introduction

Phthalates and plasticizers are synthetic chemicals that are utilized to make plastic more flexible. They are known to act as endocrine-disrupting chemicals (EDC)(Rudel et al. 2003; Hunt et al. 2009), which are ubiquitous in food and beverage containers, as well as coatings of pills, medical tubing (Green et al. 2005; R. Hauser and Calafat 2005) and plastic packaging (Muncke 2011). Phthalates and plasticizers are bound to plastic polymers by non-covalent bonds, and thus, easily leak into the environment (Pearson and Trissel 1993). The main routes of exposure to these substances are ingestion, inhalation, dermal absorption, or intravenous medication administration (R. Hauser and Calafat 2005; Meeker, Calafat, and Hauser 2009). Consequently, the vast majority of the population is exposed to these toxins on a daily basis. Low micro-molar concentrations of certain phthalates in human urine, sweat and plasma have been associated with an increased rate of miscarriages and compromised male and female fertility (Toft et al. 2012; Burdorf et al. 2011; Lovekamp and Davis 2001; Svechnikova, Svechnikov, and Söder 2007; Bloom et al. 2015; Duty et al. 2003; Russ Hauser et al. 2006; Brehm and Flaws 2019; Patel et al. 2015; W. Wang et al. 2012). One of the most commonly used phthalates is di-2-ethylhexyl phthalate (DEHP) (Y. Wang, Zhu, and Kannan 2019). In fact, 98% of the US population test positive for DEHP and its metabolites (Zota, Phillips, and Mitro 2016; Api 2001). Despite numerous reports on its toxicity, DEHP is still widely used in consumer products and in a number of medical devices, such as blood bags, infusion tubes, nasogastric tubes, peritoneal dialysis bags, and urological catheters. Patients who undergo frequent hemodialysis, catheterization or massive blood transfusions are at particular risk for DEHP toxicity and are exposed to doses as high as 168 mg/day (Kavlock et al. 2002). The amount of DEHP that will leak out of the medical device depends on the temperature and the duration of catheterization.

Several human studies have investigated the chronic effect of DEHP exposure on male fertility, primarily focusing on the links between DEHP exposure and sperm DNA damage (X.-F. Huang et al. 2012). However, limited research has been done on the acute effect of DEHP exposure on sperm capacitation, acrosome reaction, and sperm ability to successfully fertilize an egg. In this study, we used a murine model to test the acute effects of four ubiquitous environmental toxins: bisphenol A (BPA), di-2-ethylhexyl phthalate (DEHP), diethyl phthalate (DEP), and dimethyl phthalate (DMP) on sperm fertilizing ability and pre-implantation embryo development. Out of the four compounds tested, DEHP demonstrated the strongest effect on male fertility by significantly altering the profile of sperm capacitation, inhibiting acrosome reaction, and triggering excessive reactive oxygen species (ROS) production. Altogether these changes led to sperm inability to fertilize an egg. These results suggest that DEHP can directly affect sperm fertility and is therefore detrimental to male reproductive health.

## Results

### Murine embryo development is impacted by DMP, BPA, DEP, and DEHP

Exposure to phthalates could either damage sperm directly or impair embryo development after fertilization occurs. To test the susceptibility of pre-implanted embryos to DEHP, DMP, DEP, and BPA, naturally derived zygotes from 4-16-week-old female mice were harvested. The recovered zygotes were randomly divided into two groups: phthalate-free culture media and phthalate-supplemented media. The ability of the pre-implantation embryo to progress towards the blastocyst stage was recorded on day 5 post-fertilization (Figure 1). The survival rate was calculated based on the percentage of embryos that have reached the morula or blastula stage. While all four tested compounds did not affect preimplantation development at the lower concentrations (up to 2 μM), we found that at 10 μM, all four chemicals effectively prevented blastocyst formation (*p<0.05*; Figure 1A-H and Supplemental Tables 1a-1d). All controls have been performed with either vehicle control (0.1% ethanol) or drug-free media. No significant differences were observed among control conditions (Supplemental Table 2). The concentration range for phthalates was chosen based on previously reported DEHP concentrations linked to female infertility (Reddy et al. 2006).

**Figure.1.**
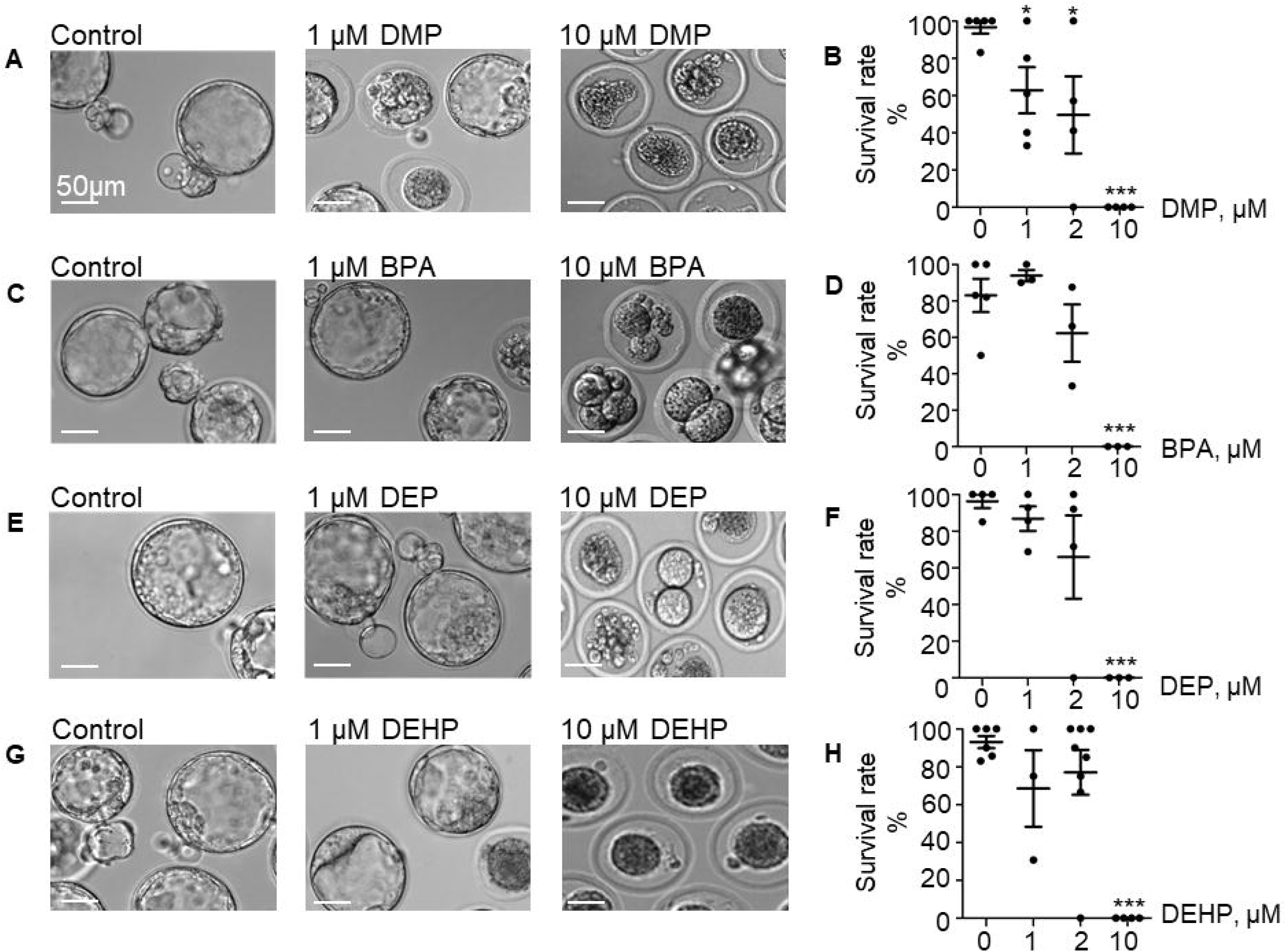
Murine embryo development is impacted by DMP, BPA, DEP and DEHP. In vitro embryo development on day 5 post fertilization. **A.** Representative Images of blastocysts previously exposed at the zygote stage to 0, 1 or 10 μM DMP, for 20 hours. Subsequent embryo culturing was done in the absences of EDCs, and the images were taken on day 5 of embryo development. Scale bar is 50 μM. **B.** The survival rate of DMP exposed zygotes was calculated based on the percentage of embryos that have reached morula or blastocyst stage. **C.** Representative images of embryos previously exposed to 0, 1 or 10 μM BPA at the same stage and duration as in (A), scale bar is 50 μM. **D.** The survival rate of BPA exposed zygotes was calculated as in (B). **E.** Representative images of embryos previously exposed to 0, 1 or 10 μM DEP at the same stage and duration as in (A), scale bar is 50 μM. **F.** The survival rate of DEP exposed zygotes was calculated as in (B). **G.** Representative images of embryos previously exposed to 0, 1 or 10 μM DEHP at the same stage and duration as in (A), scale bar is 50 μM. **H.** The survival rate of DEHP-exposed zygotes was calculated as in (B). Data are means +/− S.E.M. Asterisk indicates a statistical difference between control embryos and embryos exposed to EDCs. ** (P<0.01), *** (P<0.001).

### In vitro fertilization is affected by DEHP

To explore the direct effect of DEHP, DMP, DEP, and BPA on sperm fertilizing ability, *in vitro* fertilization (IVF) assays were conducted. Mouse sperm were capacitated in phthalate-supplemented media as described in methods by exposing sperm to three concentrations (1 μM, 2 μM, and 10 μM) of phthalates. 60-90 minutes post-exposure, sperm was introduced to healthy murine eggs. The eggs were incubated with sperm for 4 hours. The fertilization rate was calculated as described above and presented as the percentage of embryos that reached the morula or blastula stage on day 5 post-fertilization (*p<0.05*; Figure 2A-H and Supplemental Tables 3a-3d). As shown in Figure 2, all tested concentrations of DEP and DMP did not produce any effect on sperm fertilizing ability and subsequent blastocyst formation, while BPA had a minimal, but not statistically significant effect on early embryo development (Figure 2C-D). The most damaging effect to blastocyst formation was observed with DEHP (Figure 2G-H). While spermatozoa retained their fertilization potential at 1 μM, a significant decrease in embryo progression to blastulae was observed already at 2 μM (74.95 ± 5.459 % in the control vs 47.68 ± 9.68 % in 2 μM). Moreover, at 10 μM of DEHP, almost no blastocyst formation was observed (Figure 2G-H).

**Figure. 2.**
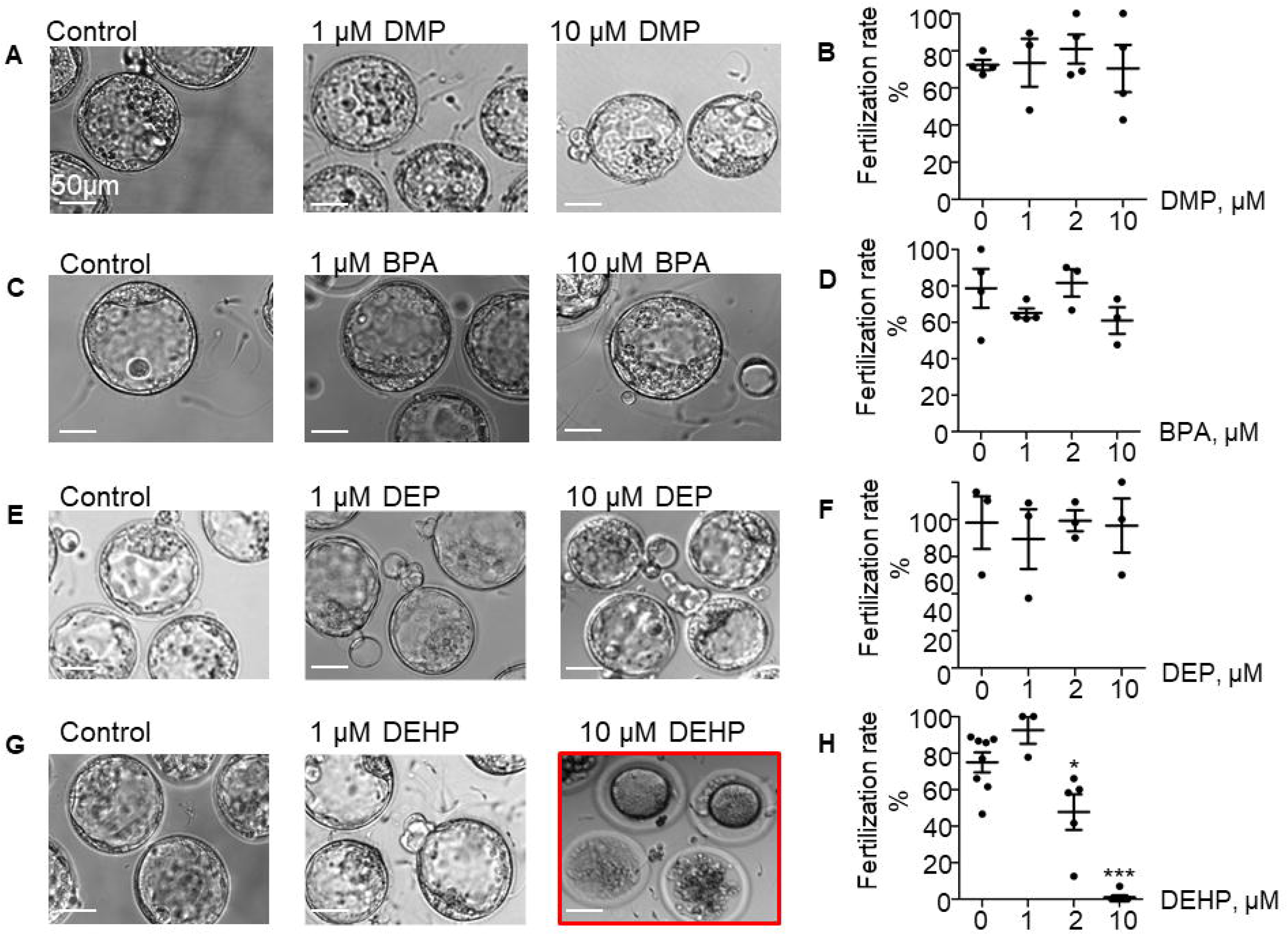
Fertilization rate of murine eggs exposed to EDCs treated spermatozoa. **A.** Representative Images of blastocysts obtained after murine eggs were introduced to sperm previously exposed to 0, 1 or 10 μM DMP. Subsequent embryo culturing was done in the absences of EDCs, and the images were taken on day 5 post insemination. Scale bar is 50 μM. **B.** Percentage of eggs that were fertilized by DMP-treated sperm and were able to reach morula or blastocyst stage. **C.** Representative Images of embryos obtained after IVF with 0, 1, 2 or 10 μM BPA-treated spermatozoa. Images were taken at the same stage as in (A), scale bar is 50 μM. **D.** The percentage of eggs fertilized by BPA-treated sperm was calculated as in (B). **E.** Representative images of embryos obtained after IVF with 0, 1, 2 or 10 μM DEP-treated sperm. Images were taken at the same stage as in (A), scale bar is 50 μM. **F.** The percentage of eggs fertilized by DEP-treated sperm was calculated as in (B). **G.** Representative Images of embryos obtained after IVF with 0, 1, 2 or 10 μM DEHP-treated sperm. Images were taken at the same stage as in (A), scale bar is 50 μM. **H.** The percentage of eggs fertilized by DEHP-treated sperm was calculated as in (B). Data are means +/− S.E.M. Asterisk indicates a statistical difference between control embryos and embryos exposed to EDCs. ** (P<0.01), *** (P<0.001).

### DEHP prevents fertilization

DEHP exposure either affects sperm’s ability to fertilize the egg or permits fertilization but inhibits zygotic division. To distinguish between these two scenarios, pronuclei formation was recorded 9 hours post *IVF*. In the presence of 10 μM DEHP, a 92.76% reduction in pronuclei formation was detected compared to untreated control (Figure 3). Values were calculated based on pooled data from three independent experiments and represent a total of 19 zygotes out of 25 eggs in the control conditions, versus 2 zygotes out of 36 eggs in the presence of DEHP (Supplemental Table 5). These results indicate that even short exposure to DEHP modifies sperm physiology making spermatozoa unable to penetrate the zona pellucida.

**Figure 3.**
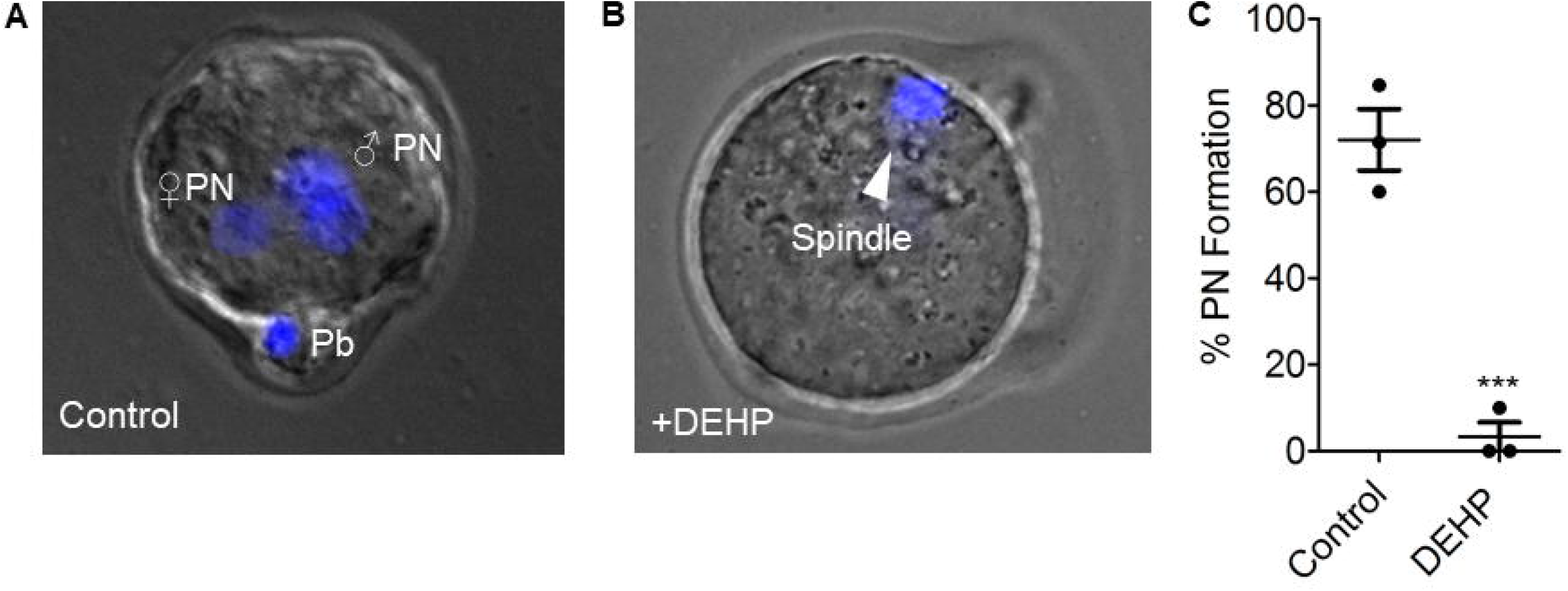
Pronuclei formation after in vitro fertilization by DEHP-treated sperm. Pronuclear formation was assessed 9 hours after IVF was performed; genomic DNA staining was done with DAPI. **A.** Representative image of a successfully fertilized egg with two pronuclei (PN) and a polar body (Pb). The egg was inseminated with sperm treated with vehicle control solution. **B.** Representative image of an unfertilized egg following an IVF with sperm that was treated with◻10 μM DEHP. The arrowhead indicates the position of the second metaphase spindle. **C.** Percentage of fertilized eggs with PN detected. Each data point represents the mean of one of the three independent experiments +/− S.E.M. *** indicates statistical significance (P<0.005). The total amount of oocytes: 25 (control), 36 (+10μM DEHP).

### Murine CatSper is not affected by DEHP

Since DEHP has demonstrated the strongest effect on sperm fertility among all tested EDCs, we further explored which sperm functions were directly affected by exposure to this phthalate. Once deposited inside the female reproductive tract, mammalian spermatozoa must undergo a final maturation step, i.e. capacitation, in order to become competent to fertilize the egg (Austin 1951; R. Yanagimachi 1994). This process results in the removal of non-covalently attached glycoproteins, depletion of cholesterol and other steroids (Davis 1981), as well as the removal of adherent seminal plasma proteins (Chang 1957). These physiological changes alter sperm membrane potential and make the cell competent to undergo a change in motility, known as hyperactivation, trigger the acrosome reaction and prepare spermatozoa for fertilization. Hyperactivation is characterized by calcium influx into the sperm flagellum via the calcium channel-CatSper (Ren et al. 2001; Carlson et al. 2003), and is defined as an asymmetrical flagellar beat that is required for penetration through the viscous luminal fluids of the female reproductive tract and the protective vestments of the egg. CatSper deficiency or its suppression by environmental toxins have been linked to male infertility (Ren et al. 2001; Qi et al. 2007; Tamburrino et al. 2014; Schiffer et al. 2014). To investigate whether murine CatSper is also affected by DEHP, we used murine sperm patch-clamp technique (Kirichok, Navarro, and Clapham 2006). As shown in Supplemental Figure 1a-1b, application of up to 100 μM of DEHP did not cause any significant changes in monovalent CatSper currents activated by a voltage ramp from −80 mV to +80 mV. This indicates that DEHP affects sperm cells through another mechanism.

### DEHP alters sperm capacitation and ROS production

Another hallmark of capacitation is the phosphorylation of sperm proteins on tyrosine residues (Salicioni et al. 2007; Pablo E Visconti 2009). Tyrosine phosphorylation occurs during the late stages of capacitation (Pablo E Visconti 2009). This allows sperm to hyperactivate, undergo the acrosome reaction and interact with the zona pellucida (Naz and Rajesh 2004). To test the effect of DEHP on sperm tyrosine phosphorylation, caudal epididymal spermatozoa were incubated in a capacitating medium containing either 10 μM DEHP or vehicle control. Subsequently, tyrosine phosphorylation was assessed via a western blot using a monoclonal anti-phosphotyrosine antibody (anti-PY) (EMD Millipore). As shown in Figure 4A-B, DEHP markedly alters sperm capacitation-associated tyrosine phosphorylation kinetics, by expediting the process within the first 60 minutes of exposure, followed by a complete reversal after 120 minutes of incubation. In addition, immunocytochemistry experiments using the same anti-PY antibody, revealed that the increased protein phosphorylation caused by DEHP is primarily localized to the mid-piece region of sperm (Figure 4C). To further assess the global changes in sperm tyrosine phosphorylation caused by DEHP, flow cytometry analysis was performed using anti-PY labeled with CF647 dye (Biotium). As shown in Figure 4D, a significant increase in overall fluorescence was observed in DEHP treated cells resulting in 1.5±0.2-fold increase in global fluorescence.

**Figure 4.**
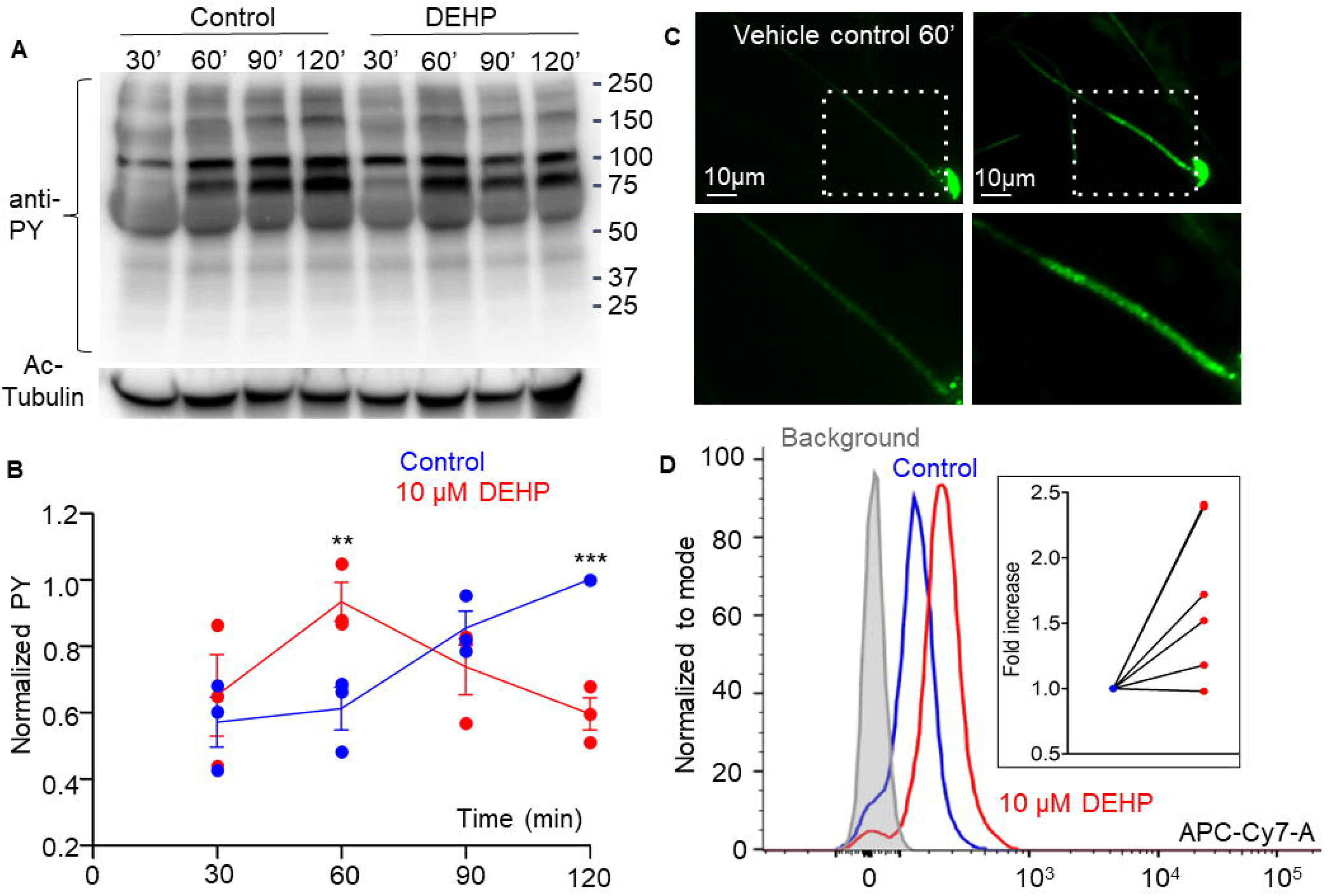
Capacitation-associated tyrosine phosphorylation of murine sperm is altered in the presence of DEHP. **A.** Representative western blot image shows the time course of protein tyrosine phosphorylation under conditions that support capacitation in the presence or absence of 10μM DEHP. DEHP was added to the media immediately before the start of capacitation process. Sperm lysates were obtained at the indicated times and subjected to SDS-PAGE immunoblotting. Tyrosine phosphorylation was detected with a monoclonal phosphor-tyrosine (PY) antibody. Acetylated-tubulin (Ac-Tubulin) was used as a loading control. **B.** Levels of relative tyrosine phosphorylation obtained as total densities extracted from (A) and normalized to the densities of the loading control. Each data point represents the mean of one of the three independent experiments. **C.** Immunofluorescent localization of phosphor-tyrosine residues as visualized by PY antibody in mouse sperm. Increased phosphorylation detected after 60 minutes of capacitation in the mid-piece region of spermatozoa in DEHP-treatment group (right panel) as compared to control untreated spermatozoa (left upper panel). Lower panels represent insets from the corresponding region of interests indicated on the upper panels by dashed rectangular. **D.** A representative flow cytometry data showing an increase in global tyrosine phosphorylation in 10μM DEHP-treated spermatozoa (red) at 60 minutes of capacitation compared to the vehicle control (blue). Tyrosine phosphoproteins were detected using a CF 647 dye conjugated to an anti-PY (mAb). Inset: fold increase in mean fluorescent intensity normalized to mode as detected by the flow cytometer compared to control conditions. Data are means +/− S.E.M. ** indicates statistical significance (P<0.01) between control spermatozoa and spermatozoa exposed to 10μM DEHP. *** indicates statistical significance (P<0.001)

The mid-piece region of sperm flagellum harbors mitochondria-an organelle known to generate reactive oxygen species (ROS). Interestingly, DEHP increases ROS generation in various cells and tissues, including hepatocytes, adipocytes, and testis (Y. Huang et al. 2019; Kasahara et al. 2002; Schaedlich et al. 2018). However, DEHP’s ability to alter sperm ROS production has not been studied. To detect ROS production, a chemiluminescence assay was employed, a commonly described technique to detect ROS in semen (Ochsendorf et al. n.d.; Williams and Ford 2005; Agarwal et al. 2008). The levels of ROS production were assessed in caudal sperm capacitated in the presence or absence of 10 μM and 100 μM DEHP after 60 minutes of exposure. A significant increase in ROS production was detected in all treated samples in comparison to vehicle-treated controls (Figure 5A). These results indicate that in addition to changes in tyrosine phosphorylation, DEHP triggers an increase in ROS production in sperm. Since excessive ROS production is known to be cytotoxic to sperm, exposure to DEHP may lead to impaired sperm fertilizing capacity.

**Figure 5.**
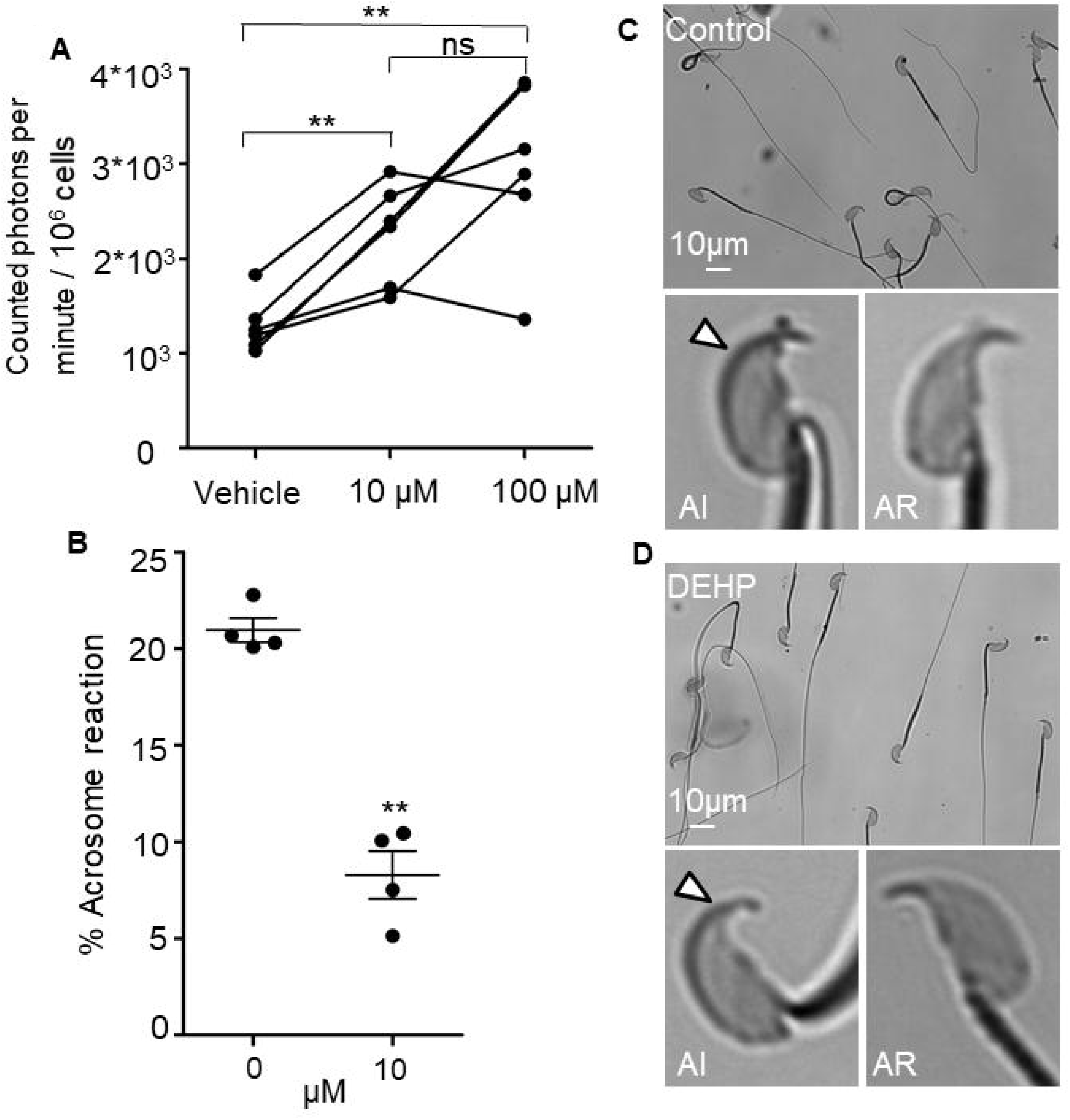
DEHP increases ROS production and decreases spontaneous acrosome reaction in capacitated sperm cells. **A.** Luminol-dependent chemiluminescence assay showed increased rates of ROS production in sperm treated with 10 and 100 μM DEHP in comparison to the vehicle control (0 μM DEHP). Connected dots represent spermatozoa extracted from the same mouse divided into 3 aliquots and subjected to the different conditions (n=6). A minimum of 1.35*10^6^ sperm cells/mL were used per condition. **B.** Percentage of spontaneous acrosome-reaction in capacitated spermatozoa with or without 10μM DEHP. Each data point represents the mean of one of the four independent experiments. A minimum of 500 cells were evaluated per experiment. **C.** Acrosome-reacted (AR) and Acrosome-intact (AI) spermatozoa in the control condition. **D.** AR and AI spermatozoa in 10μM DEHP condition. Arrowheads point at intact acrosome. Data are means +/− S.E.M. ** indicates statistical significance (P<0.01) between control spermatozoa and spermatozoa exposed to 10μM DEHP.

### The acrosome reaction in capacitated spermatozoa is inhibited by DEHP

The acrosome reaction is the fusion of sperm plasma membrane with the outer acrosomal membrane and is vital for fertilization as only acrosome-reacted spermatozoa are able to fuse with the egg (Avella and Dean 2011). According to previous reports, increased levels of ROS production result in excessive peroxidation of the sperm acrosomal membrane (Zalata et al. 2004), impairing acrosomal exocytosis and sperm-egg fusion (R J Aitken and Clarkson 1987; Ichikawa et al. 1999; Griveau, Renard, and Le Lannou 1995). While there is a debate over the physiological triggers for the acrosome reaction and the exact site of acrosomal exocytosis, it is well accepted that the acrosome reaction is required for sperm fertility. Therefore, we have explored whether DEHP can alter spontaneous sperm acrosome reaction. Indeed, incubation with 10 μM DEHP reduced the percentage of sperm that underwent the spontaneous acrosome reaction in capacitated sperm from 20.95 ±0.62 % in control to 8.28 ±1.24 % in the presence of DEHP (Figure 5B-D). Together, these results indicate that DEHP exposure triggers excessive ROS production in sperm, as well as resulting in altered tyrosine phosphorylation and acrosome reaction. Consequently, these changes negatively affect sperm fertility.

## Discussion

Exposure to EDCs poses a significant risk to reproductive health and fetal development (Brehm et al. 2018; Hannon, Niermann, and Flaws 2016). While it is known that DEHP impairs sperm motility and chromatin DNA stability (Sumner et al. 2019; Pant et al. 2011), its mechanism of action and direct effect on sperm fertilizing capacity have not been elucidated. In this study, we aimed to determine the acute effect of DEHP on sperm fertility. Upon absorption, DEHP is distributed throughout the body by the circulatory system. The majority of DEHP is quickly hydrolyzed by the liver into various metabolites, which have been linked to altered fertility and DNA damage in sperm (R. Hauser et al. 2007). However, a portion of DEHP is stored un-metabolized in the adipose tissue-which acts as a reservoir for lipophilic EDCs (Tanaka et al. 1975; Regnier and Sargis 2014). Hormonal and neuronal signals regulating the fat tissue can trigger an abrupt release of lipophilic EDCs to the systemic circulation (Regnier and Sargis 2014). There are two main ways in which sperm can encounter un-metabolized DEHP: via abrupt release from adipose tissue or through release from medical devices, such as a urological catheter-making unmetabolized DEHP a prominent threat to sperm fertility.

DEHP exposure has been previously linked to increased ROS production in somatic cells and oocytes (Wu et al. 2014; Tripathi et al. 2019; Kim et al. 2013; Ambruosi et al. 2011). However, its impact on ROS over-production in sperm was not elucidated. Here we show that DEHP exposure triggers excessive ROS generation, leading to oxidative damage and ultimately sperm infertility.

While minor ROS generation naturally takes place during early sperm capacitation, a maturation step spermatozoa undergo to acquire fertilizing capacity, this process must be tightly regulated. Mild ROS production triggers an increase in intracellular cAMP, resulting in the activation of Protein Kinase A (PKA). PKA, in turn, carries out a series of controlled tyrosine phosphorylation events in a time-dependent manner. (R. J. Aitken et al. 1995; Leclerc, De Lamirande, and Gagnon 1997). Interestingly, certain EDCs such as BPA have been shown to up-regulate PKA’s activity leading to an altered phosphorylation pattern downstream of PKA (Rahman et al. 2015). Since mature spermatozoa are transcriptionally and translationally silent cells, post-translational modifications such as protein tyrosine phosphorylation play an essential role in their maturation process and ability to fertilize an egg (Naz and Rajesh 2004). Unwarranted ROS production leads to over-phosphorylation which significantly alters the maturation process of sperm (Donà et al. 2011; Villegas et al. 2003). Moreover, an excess of ROS is cytotoxic to sperm due to their high content of polyunsaturated fatty acids in the plasma membrane and their limited antioxidant capacity (R. J. Aitken and Clarkson 1987; Alvarez and Storey 1982; Jones, Mann, and Sherins 1979).

DEHP-treated sperm cells had altered capacitation with aberrantly fast tyrosine phosphorylation within the first 60 minutes of capacitation. This differs from the gradual increasing protein phosphorylation pattern that was observed in the control condition and previously reported in the literature (P. E. Visconti et al. 1995; Piehler et al. 2006). The detected increase in tyrosine phosphorylation was localized primarily to the mid-piece region of sperm, the flagellar compartment where mitochondria are located. Mitochondrial respiration produces a significant amount of ROS, this process, if unregulated, can damage sperm genomic DNA, lipid and protein structures, and subsequently impair sperm integrity and fertility. Previous reports show that DEHP exposed oocytes and somatic cells produce an excessive amount of ROS via mitochondrial-derived ROS (Rosado-Berrios, Vélez, and Zayas 2011; Roth 2018; Wu et al. 2014). However, it has not been shown that DEHP affects spermatozoa in a similar manner. In fact, several EDCs were suggested to affect sperm fertility via CatSper-related mechanism (Schiffer et al. 2014). Here, we show that while murine CatSper was not sensitive to DEHP exposure, this phthalate indeed triggers excessive ROS production and subsequently impairs sperm fertility.

An additional effect of excessive oxidative stress on spermatozoa is lipid peroxidation. Spermatozoa are extremely susceptible to lipid peroxidation due to their high concentration of polyunsaturated fatty acids (R. John Aitken et al. 2006; Wathes, Abayasekara, and Aitken 2007; R J Aitken and Clarkson 1987; Alvarez and Storey 1982; Jones, Mann, and Sherins 1979). Alteration of the lipid structure due to peroxidation in sperm leads to a decrease in membrane fluidity (Chen et al. 2013; Cocuzza et al. 2007; Sikka 2001; Zalata et al. 2004) causing decreased motility and altered acrosome reaction (Zalata et al. 2004; R J Aitken and Clarkson 1987; Ichikawa et al. 1999; Griveau, Renard, and Le Lannou 1995). The acrosome reaction is an important step during fertilization. Murine sperm begin to undergo the acrosome reaction in the upper isthmus (La Spina et al. 2016), the part of the oviduct that connects the uterine with the ampulla. Thus, mouse fertile sperm are acrosome-reacted prior to reaching the ampulla, the site of fertilization and before encountering the eggs (Jin et al. 2011). In fact, most acrosome-intact spermatozoa are unable to fertilize the egg and swim away from the zona pellucida (Jin et al. 2011). Thus, sperm ability to undergo the AR at the end of capacitation is highly important for sperm fertility (Ryuzo Yanagimachi 2011). Here we find that DEHP significantly inhibits the acrosome reaction in capacitated sperm. As a result, DEHP-exposed sperm were largely acrosome-intact and therefore unable to fertilize murine eggs. This explains the reduced levels of fertilization that were observed in IVF and the absence of pronuclei formation. These results indicate that, in addition to its chronic impact on reproductive potential, DEHP also imposes acute damage to sperm by affecting its ability to fertilize and thereby represent a risk to male fertility.

## Figure legends

**Supplementary Figure 1.**
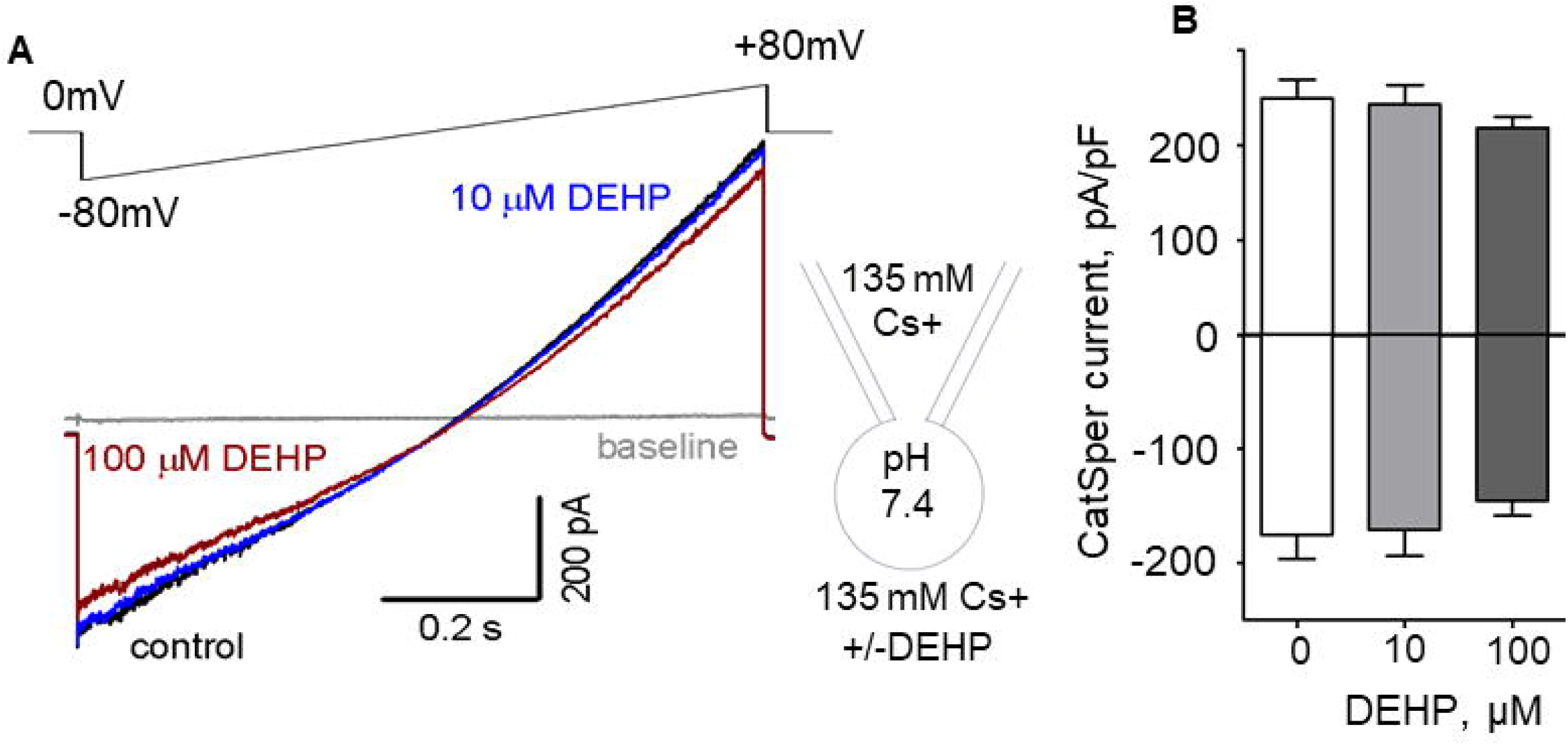
Murine CatSper is not affected by DEHP. **A.** Representative monovalent whole-cell CatSper currents (*I_CatSper_*) recorded from a murine spermatozoon in the absence (black) and presence of 10 μM (blue) and 100 μM DEHP (red). *I_CatSper_* were activated by a voltage ramp from −80 mV to +80 mV from a holding potential of 0 mV. Voltage protocol is shown above the currents. The panel on the right shows the main conducting ion of the pipette and bath solutions. **B.** Averaged *I_CatSper_* densities recorded from murine epididymal spermatozoa in the absence and presence of DEHP. Data are means +/− S.E.M. An average of 3 independent experiments is shown.

**Supplementary Figure 2.**
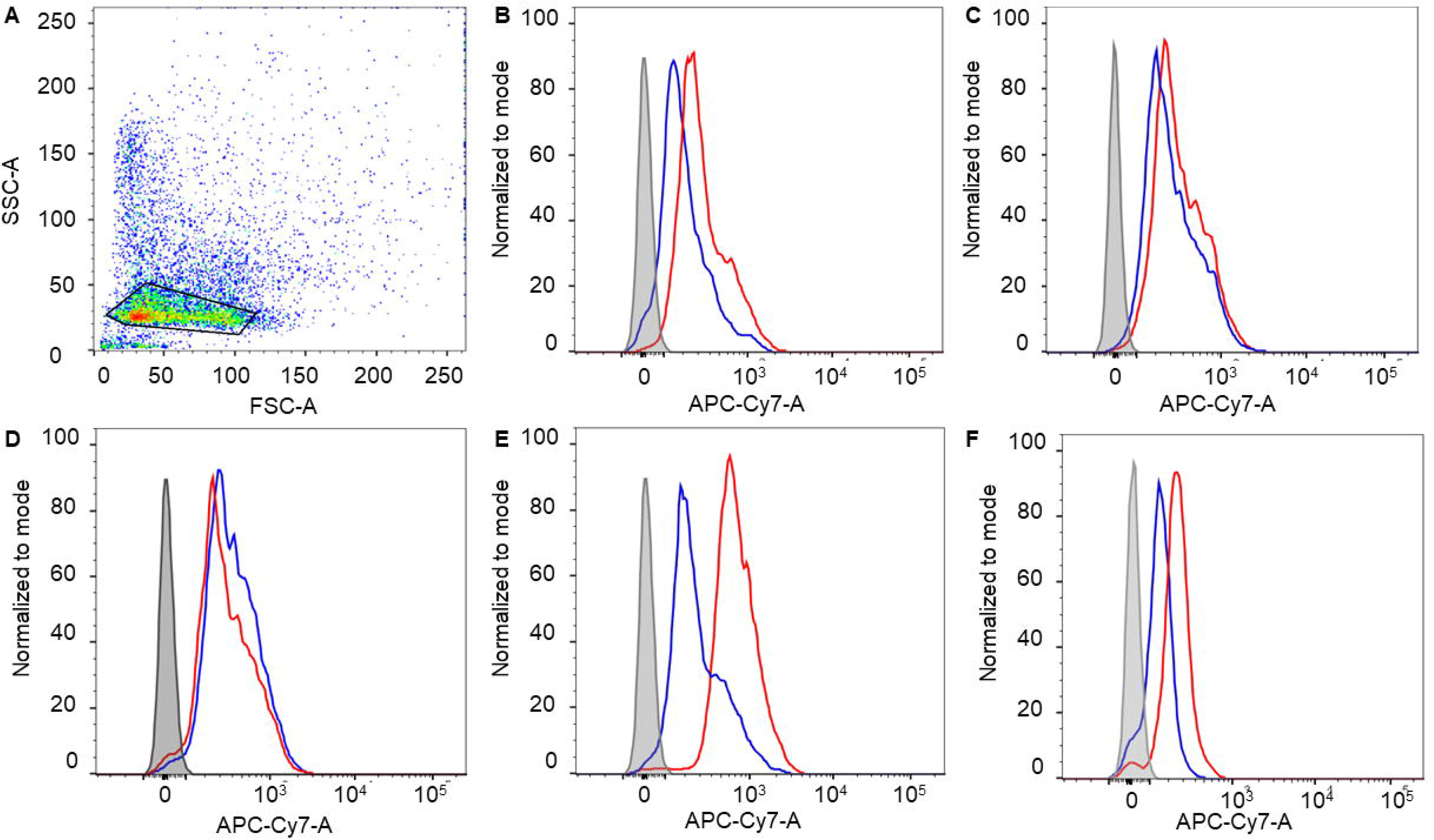
Representative dot plot of side (SSC-A) versus forward (FSC-A) scatter showing flow-cytometry data obtained for sperm. **A.** The region of interest demarcated by solid lines was selected to eliminate cellular debris. **B-F.** Representative flow cytometry histograms from five independent experiments. Mean fluorescence intensities (MFI) normalized to mode show an increase in global tyrosine phosphorylation in 10μM DEHP treated spermatozoa (red) at 60 minutes of capacitation compared to the vehicle control (blue). The background fluorescence detected in unstained spermatozoa is shown in grey. Tyrosine phosphoproteins were detected using a CF 647 dye conjugated to anti-PY (monoclonal antibody).

**Supplementary Tables 1A-1D.**
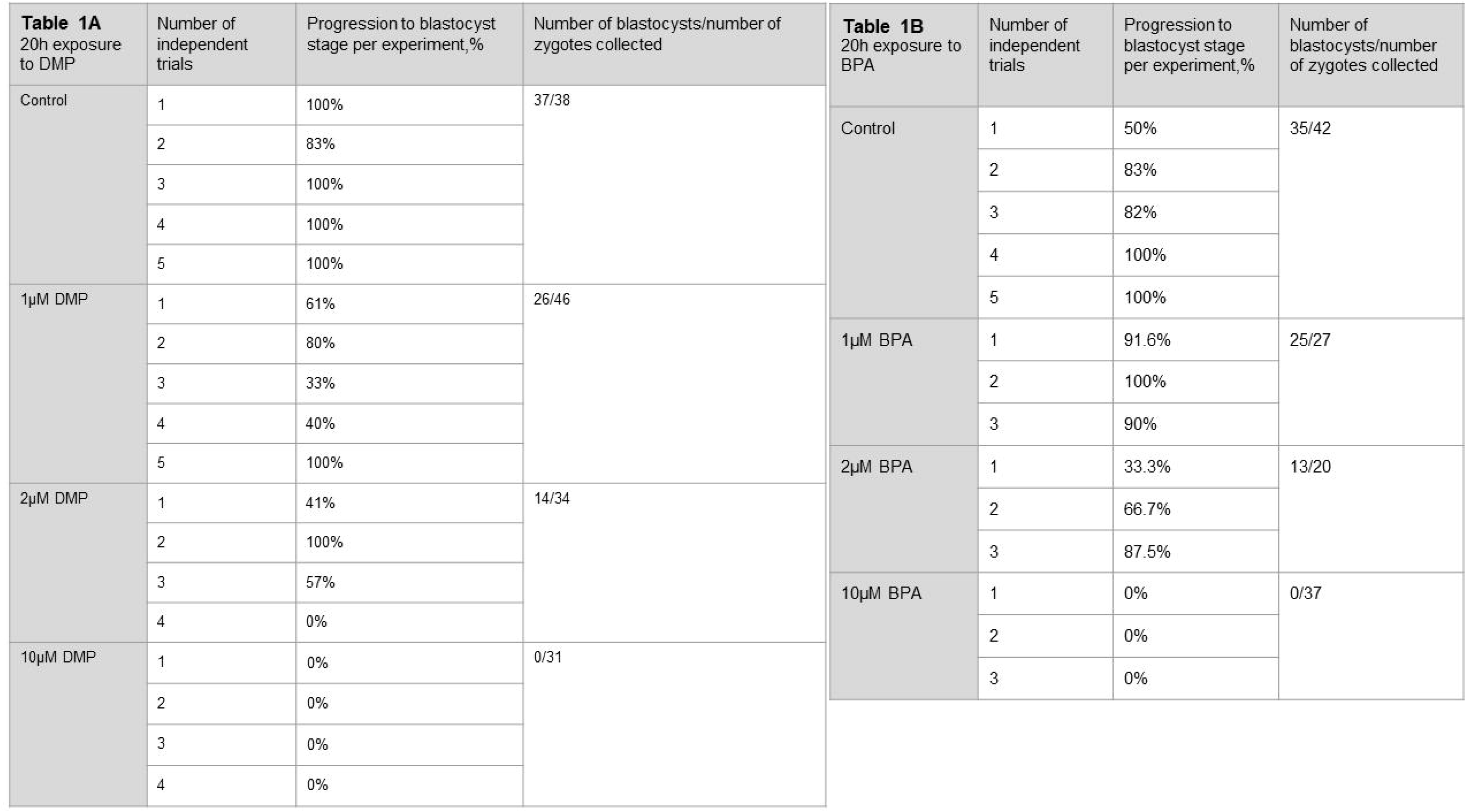
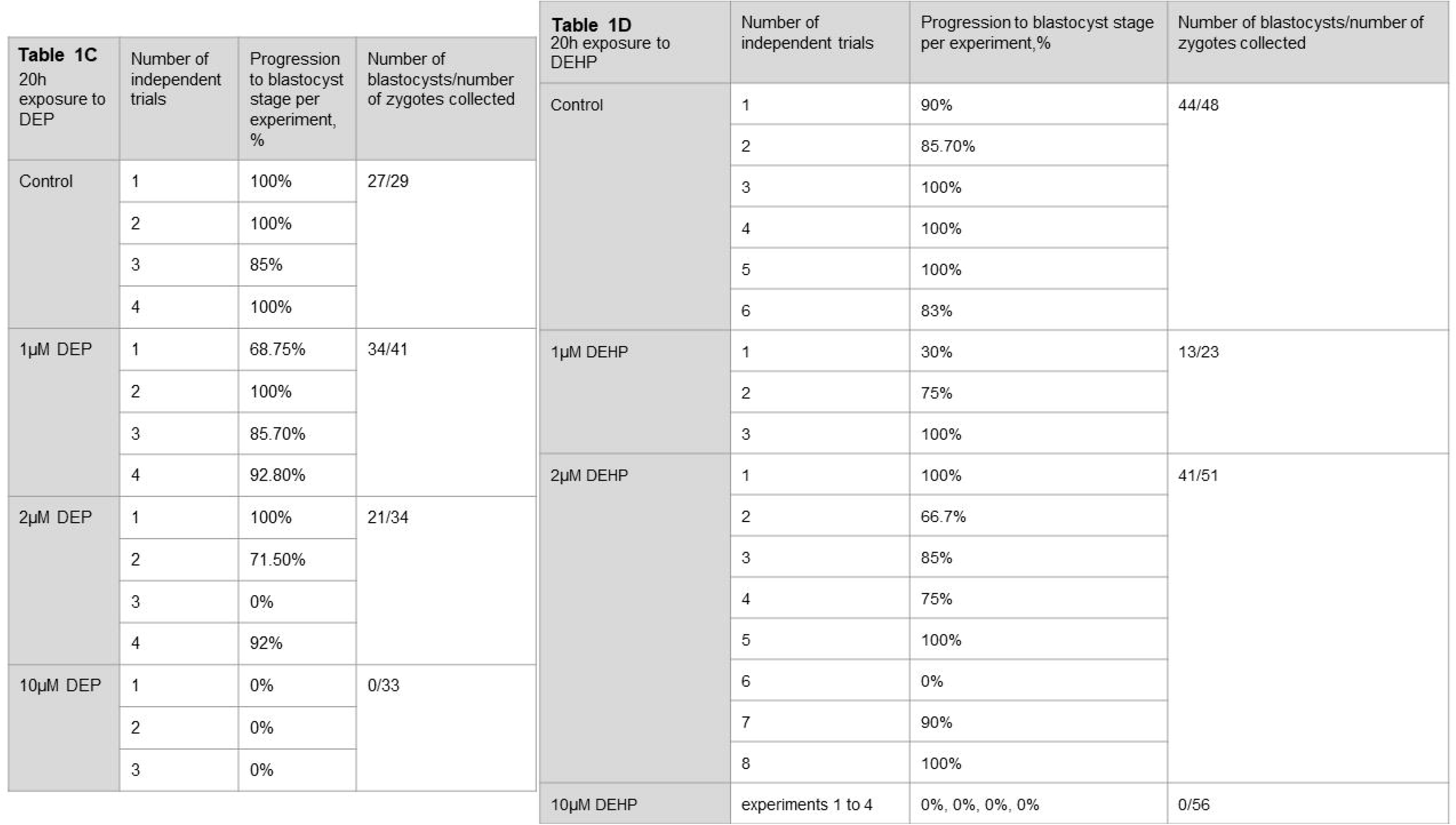
Murine embryo development after 20h exposure to DMP, BPA, DEP or DEHP. Embryo development was assessed on day 5 post fertilization. The column “progression to blastocyst stage per experiment, %” represents the percentage of embryos that reached blastocyst or morula stage. This number was calculated by dividing the number of all embryos that reached blastocyst or morula stage to the number of all collected zygotes per each experiment. Zygotes were obtained from naturally mated super-ovulated females. Each condition was assessed by 3-8 independent experiments. **A.** Embryo development after 20 hours exposure to DMP at 0, 1, 2 and 10μM and subsequent embryo culture in DMP-free media. **B.** Embryo development after 20 hours exposure to the indicated concentration of BPA and subsequent culture in BPA-free media. **C**. Embryo development after 20 hours exposure to the indicated concentration of DEP and subsequent culture in DEP-free media. **D.** Embryo development after 20 hours exposure to the indicated concentration of DEHP and subsequent culture in DEHP-free media.

**Supplementary Table 2.** The development of murine zygotes isolated from naturally mated super-ovulated females and after their exposure to 0.1% ethanol for 20 hours in the culture media. Embryo development was assessed on day 5 post fertilization and represents the percentage of embryos that reached blastocyst or morula stage. This number was calculated by dividing the number of all embryos that reached blastocyst or morula stage by the number of all collected zygotes per experiment.

| 20h exposure to Vehicle Control, 1μL/mL | Number of independent trials | Progression to blastocyst stage per experiment, % | Number of blastocysts/number of zygotes collected |
| --- | --- | --- | --- |
| Vehicle Control, 1μL/mL | 1 | 100% | 16/17 |
|  | 2 | 83% |  |
|  | 3 | 100% |  |

**Supplementary Tables 3A-3D.** Development of in vitro fertilized mouse embryos obtained after murine eggs were introduced to the sperm previously exposed to DMP, BPA, DEP or DEHP for 60-90 minutes. Embryo development was assessed on day 5 post fertilization. The “Progression to the blastocyst stage per experiment, %” column represent the percentage of embryos that reached blastocyst or morula stage. This number was calculated by dividing the number of all embryos that reached blastocyst or morula stage to the number of all collected and inseminated eggs per each experiment. Each condition was assessed by 3-5 independent experiments. **A.** *In vitro* embryo development after eggs insemination with 0, 1, 2 or 10μM DMP-treated sperm **B.** *In vitro* embryo development after eggs were inseminated with spermatozoa previously exposed to the indicated concentration of BPA. **C**. *In vitro* embryo development after eggs were inseminated with the spermatozoa treated with corresponding concentrations of DEP. **D.** *In vitro* embryo development after murine egg were inseminated with spermatozoa treated with corresponding concentrations of DEHP.

| **Table 3A**<br>IVF; DMP | Number of independent trials | Progression to blastocyst stage per experiment, % | Number of blastocysts/number of eggs collected |
| --- | --- | --- | --- |
| Control | 1 | 71.4% | 26/36 |
|  | 2 | 71% |  |
|  | 3 | 80% |  |
|  | 4 | 67% |  |
| 1µM DMP | 1 | 48% | 30/42 |
|  | 2 | 89.5% |  |
|  | 3 | 83% |  |
| 2µM DMP | 1 | 69% | 28/36 |
|  | 2 | 100% |  |
|  | 3 | 87.5% |  |
|  | 4 | 67% |  |
| 10µM DMP | 1 | 57% | 21/30 |
|  | 2 | 43% |  |
|  | 3 | 82% |  |
|  | 4 | 100% |  |

| **Table 3 B**<br>IVF; BPA | Number of independent trials | Progression to blastocyst stage per experiment, % | Number of blastocysts/number of eggs collected |
| --- | --- | --- | --- |
| Control | 1 | 100% | 24/28 |
|  | 2 | 88% |  |
|  | 3 | 77% |  |
|  | 4 | 50% |  |
| 1µM BPA | 1 | 62% | 56/88 |
|  | 2 | 73% |  |
|  | 3 | 63% |  |
|  | 4 | 63% |  |
| 2µM BPA | 1 | 90% | 39/47 |
|  | 2 | 88% |  |
|  | 3 | 67% |  |
| 10µM BPA | 1 | 73% | 13/19 |
|  | 2 | 63% |  |
|  | 3 | 48% |  |

| Table 3C<br>IVF; DEP | Number of independent trials | Progression to the blastocysts stage per experiment, % | Number of blastocysts/number of eggs collected |
| --- | --- | --- | --- |
| Control | 1 | 90% | 28/31 |
|  | 2 | 94.7% |  |
|  | 3 | 50% |  |
| 1µM DEP | 1 | 88.8% | 20/28 |
|  | 2 | 82% |  |
|  | 3 | 38% |  |
| 2µM DEP | 1 | 89% | 35/42 |
|  | 2 | 78% |  |
|  | 3 | 70% |  |
| 10µM DEP | 1 | 80% | 18/20 |
|  | 2 | 100% |  |
|  | 3 | 50% |  |

| Table 3D<br>IVF; DEHP | Number of independent trials | Progression to the blastocyst stage per experiment, % | Number of blastocysts/number of eggs collected |
| --- | --- | --- | --- |
| Control | 1 | 86% | 72/92 |
|  | 2 | 87% |  |
|  | 3 | 89% |  |
|  | 4 | 88% |  |
|  | 5 | 77% |  |
|  | 6 | 65% |  |
|  | 7 | 47% |  |
|  | 8 | 67% |  |
| 1µM DEHP | 1 | 77% | 16/18 |
|  | 2 | 100% |  |
|  | 3 | 100% |  |
| 2µM DEHP | 1 | 58% | 25/51 |
|  | 2 | 60% |  |
|  | 3 | 13% |  |
|  | 4 | 66% |  |
|  | 5 | 42% |  |
| 10µM DEHP | experiments 1 to 7 | 0%, 0%, 0%, 0%, 0%, 0%, 0%, 7% | 1/63 |

**Supplementary Table 4.** Development of embryos derived from the murine eggs that were subjected to in vitro fertilization (IVF) with murine sperm previously exposed to 0.1% ethanol for 60-90 minutes. Embryo development was assessed on the day 5 post IVF and represents the percentage of embryos that reached blastocyst or morula stage. This number was calculated by dividing the number of all embryos that reached blastocyst or morula stage to the number of all collected eggs per each independent experiment.

| 20h exposure to Vehicle Control, 1μL/mL | Number of independent trials | Progression to blastocyst stage per experiment, % | Number of blastocysts/number of zygotes collected |
| --- | --- | --- | --- |
|  | 1 | 100% | 28/37 |
|  | 2 | 80% |  |
|  | 3 | 100% |  |
|  | 4 | 50% |  |

**Supplementary Table 5.** Effect of 10μM DEHP on fertilization and pronuclei formation. Pronucleus (PN) formation was assessed 9 hours after egg insemination with sperm cells previously exposed to 10μM DEHP. Three independent experiments were carried out.

| **Table 5**<br>**PN**<br>**formation** | Number of<br>independent trials | Progression to blastocyst<br>stage per experiment, % | Number of<br>blastocysts/number of<br>zygotes collected |
| --- | --- | --- | --- |
| Control | 1 | 84.6% | 21/25 |
|  | 2 | 71.4% |  |
|  | 3 | 60% |  |
| DEHP<br>10µM | 1 | 0% | 2/36 |
|  | 2 | 0% |  |
|  | 3 | 10% |  |

**Supplementary Table 6.** Assessment of the global tyrosine phosphorylation by flow cytometry. Tyrosine phosphoproteins were detected by flow cytometry using a CF 647 dye conjugated to anti-PY (mAb). 5 independent experiments were carried out. Sperm concentrations were normalized between all conditions for each experiment. Values are mean fluorescence intensity (MFI) detected in the APC-Cy7-A channel normalized to mode. A ratio of the normalized detected MFI in the 10μM DEHP-treated sperm cells vs the vehicle control was used to calculate the fold increase in global fluorescence.

| **Table 6**<br>**Flow**<br>**cytometry**<br>Mean<br>Fluorescence<br>Intensity<br>(MFI)<br>APC-Cy7-A<br>normalized to<br>mode | Number of<br>independent<br>trials | Control<br>MFI | DEHP<br>MFI | MFI<br>Ratio<br>DEHP/Control |
| --- | --- | --- | --- | --- |
|  | 1 | 151 | 260 | 1.72 |
|  | 2 | 374 | 442 | 1.18 |
|  | 3 | 265 | 402 | 1.52 |
|  | 4 | 523 | 514 | 0.98 |
|  | 5 | 358 | 859 | 2.39 |

**Supplementary Table 7.** Detection of reactive oxygen species (ROS) in murine spermatozoa treated with DEHP compared with vehicle control. The luminol-dependent chemiluminescence assay was used to detect ROS production. Six independent experiments were carried out. A minimum of 1.35*10^6^ sperm cells/mL were used per condition. ROS production was expressed as counted photons per minute (CPM)/ 10^6^. Each row represents an individual experiment. For each experiment, sperm concentrations were normalized between all conditions.

| Mouse # | cell/mL in vehicle | Vehicle CPM | CPM/ 10<sup>6</sup> sperm vehicle | cell/mL in 10μM | 10μM CPM | CPM/ 10<sup>6</sup> sperm 10μM | cell/mL in 100μM | 100μM CPM | CPM/ 10<sup>6</sup> sperm 100μM |
| --- | --- | --- | --- | --- | --- | --- | --- | --- | --- |
| 1 | 2.57*10<sup>6</sup> | 4694 | 1826.459 | 2.17*10<sup>6</sup> | 6321.666 | 2913.210 | 2.525*10<sup>6</sup> | 6750.333 | 2673.399 |
| 2 | 1.4*10<sup>6</sup> | 1520 | 1085.714 | 1.375*10<sup>6</sup> | 3214 | 2337.454 | 1.4*10<sup>6</sup> | 5345.333 | 3818.095 |
| 3 | 1.425*10<sup>6</sup> | 1777.333 | 1247.251 | 1.9*10<sup>6</sup> | 3216 | 1692.631 | 2.775*10<sup>6</sup> | 3769 | 1358.198 |
| 4 | 1.35*10<sup>6</sup> | 1612 | 1194.074 | 2.1*10<sup>6</sup> | 3329.666 | 1585.555 | 1.775*10<sup>6</sup> | 5127.666 | 2888.826 |
| 5 | 2.375*10<sup>6</sup> | 3232 | 1360.842 | 2.25*10<sup>6</sup> | 5983.666 | 2659.407 | 1.675*10<sup>6</sup> | 5278 | 3151.044 |
| 6 | 1.575*10<sup>6</sup> | 1619.666 | 1028.359 | 1.475*10<sup>6</sup> | 3526.333 | 2390.734 | 1.225*10<sup>6</sup> | 4722.333 | 3854.965 |

## Methods

### Ethical statements and animal care

All experiments were performed in accordance with NIH Guidelines for Animal Research and approved by UC Berkeley Animal Care and Use Committee (AUP 2015-07-7742), with every effort made to minimize suffering for the animals. C57BL6 mice were purchased from Jackson Laboratory, Bar Harbor, Maine or Harlan Laboratories (Indianapolis, IN). The mice were kept in a room in a room with controlled light (14 hours light, 10 hours darkness) and temperature (23 ± 0.5°C); 50–60% humidity. The mice were fed a standard chow diet (PicoLab Rodent diet 20, LabDiet, 5053) and hyper-chlorinated water *ad libitum*.

BPA, DEHP, DMP, and DEP (Sigma-Aldrich, St. Louis, MO) were dissolved in ethanol (Sigma-Aldrich). Phthalates were used at a final concentration of 1 μM, 2 μM, and 10 μM. Vehicle controls were performed at the highest concentration.

### Embryo Collection from Natural Mating

Superovulation was induced in 4- to 16 week-old female mice by i.p injection of 5 IU of pregnant mare serum gonadotropin (5 IU, PMSG; EMD Millipore) at 14:30 and 48 hours later with 5 IU of human chorionic gonadotropin (5 IU, hCG; EMD Millipore). At the time of hCG injection, each female mouse was placed in an individual cage with one proven breeder (3-10 months old). The following morning, female mice were inspected for vaginal plugs. 20h after hCG administration, embryos were dissected out from the oviducts. The isolated oocyte-cumulus complexes were placed in pre-warmed 50 μL droplet of Hyaluronidase (80 IU/mL) (LifeGlobal) and were gently pipetted up and down repeatedly in a fine glass pipette until the oocytes were partially denuded. The oocytes were then transferred to pre-warmed M2 media (Zenith Biotech) supplemented with 4 mg/mL BSA (Sigma-Aldrich) and washed in 4-5 droplets until all the corona cells were removed. Zygotes from each individual mouse were randomly allocated to different culture conditions for 20h of incubation.

Since the tested EDCs show low water solubility and high oil solubility, the standard culture of embryos under oil could not be employed. Thus, we cultured the embryos in 500 μL KSOM (Zenith biotech) +/− EDCs at different concentrations for 20h in 4-well dishes ((Nunc™), Sigma-Aldrich) without oil. At the end of the 20h incubation, the two cell embryos were briefly washed and allocated to culture dishes, containing 10 μL droplets of KSOM (Zenith biotech) supplemented with 1 mg/mL BSA overlaid with embryo-suitable light mineral oil (Millipore) in 5% CO_2_ and 37°C. Successful development was considered as morula or blastocyst – the final stage of embryonic development before implantation.

### In Vitro Fertilization

To investigate the influence of EDCs exposure on spermatozoa’s ability to fertilize eggs, *IVF* experiments were conducted. Eggs were recovered from 4 to 16-week-old female mice by superovulation as described above. 13 hours after hCG injection, the female mice were euthanized, and the oviducts were dissected out. The oocyte-cumulus complexes were isolated and incubated in HTF medium (Embryomax, Specialty Media, Millipore), 5% CO_2_, 37°C for 30 minutes prior to insemination.

Sperm was obtained from a mature C57BL male mouse of proven fertility just before egg harvest. Spermatozoa were recovered by removing the caudae epididymis and placing each cauda separately in a petri dish containing pre-warmed HTF with or without EDCs. The tissue was cut five to six times, and sperm was allowed to swim out into the medium for 20-30 min. The cauda was then removed, and the resultant sperm suspension was left for an additional 30-60 minutes in the media at 37°C in 5% CO2 to capacitate. Total time of capacitation was 60-90min.

Four well plates (Nunc™) were used for fertilization. Each well was filled with 700 μL HTF and there, eggs were mixed with capacitated spermatozoa to a final concentration of 2.5×10^5^ cells/mL. Dishes were placed in the incubator and maintained at 37°C in 5% CO2 for 4h. After that time, eggs were washed to remove excess sperm and then cultured in 10 μL droplets of KSOM (Zenith biotech) supplemented with 1 mg/mL BSA (Sigma-Aldrich) and overlaid with embryo-tested light mineral oil (Millipore) in 5% CO_2_ and 37°C. Fertility was considered as the percentage of morula or blastocyst embryos produced by *IVF* on day five post insemination.

### Electrophysiological Analysis

Sperm was collected as described previously (Wennemuth et al. 2003). Gigaohm seals between the patch pipette and mouse spermatozoa were formed at the cytoplasmic droplet. Seals were formed in HS solution comprising the following (in mM): 130 NaCl, 5 KCl, 1 MgSO_4_, 2 CaCl_2_, 5 glucose, 1 sodium pyruvate, 10 lactic acid, 20 HEPES, pH 7.4 adjusted with NaOH. Transition into the whole-cell mode was performed by applying short (1◻ms) 499–611◻mV voltage pulses, combined with light suction. Access resistance was 15- 25◻MΩ. Cells were stimulated every 5◻s. Data were sampled at 2–5◻kHz and filtered at 1◻kHz. Pipettes (15–20◻MΩ) for whole-cell patch-clamp recordings of monovalent CatSper currents were filled with the following (in mM): 130 Cs-methane sulfonate, 70 HEPES/MES, 3 EGTA, 2 EDTA, 0.5 Tris-HCl, pH 7.4 adjusted with CsOH. Bath divalent-free solution for recording of monovalent CatSper currents contained the following (in mM): 140 Cs-methane sulfonate, 40 HEPES/MES, 1 EDTA, pH 7.4 adjusted with CsOH. HS solution was used to record baseline current while measuring monovalent CatSper currents. 1 μL/mL EtOH (vehicle control), 10 μM or 100 μm DEHP were added to the bath solution right before electrophysiology experiments. CaCl_2_ was added to this solution in accordance with WinMAXC version 2.05 (C. Patton, Stanford University) to obtain the required free Ca^2+^ concentration.

### Capacitation of Spermatozoa in the Presence of 10 μM DEHP to Test the Level of Tyrosine Phosphorylation

Spermatozoa were recovered by removing the cauda epididymis and placing it into a Petri dish containing HTF with either 1 μL/mL ethanol or 10 μM DEHP. The tissue was cut five to six times, and sperm was allowed to swim out for 15-20 minutes at 37°C in 5% CO2. The cauda was then removed, and spermatozoa suspension was further incubated at 37°C in 5% CO2. Sperm samples were collected after 30, 60, 90, and 120 minutes of capacitation and placed into a clean tube. After each collection, the sample was centrifuged at 21000 × *g* for 1 minute. The supernatant was discarded, and the cellular pellet was re-suspended in 25 μL of 2x Laemmli sample buffer (BioRad). The sample was then boiled for 5 minutes at 95°C and centrifuged at 21000 × *g* for 1 minute. Supernatants were transferred to clean tubes, β-mercapto-ethanol was added (to a final concentration of 2.5%), and the sample was heated again to 95°C for 1 min. 20 μL of the total crude cell lysate from each sample was loaded onto a 4–20% gradient Tris-HCl Criterion SDS-PAGE (BioRad). After transfer to polyvinylidene fluoride membrane, blots were blocked in 0.1% PBS-Tween20 (Fisher Scientific) with 3% IgG-free BSA for one hour and incubated with anti-phosphotyrosine antibody, clone 4G10 (Millipore, 05-321) at a dilution of 1:2000 in 1% blocking solution overnight at +4°C. The membrane was then washed three times in PBST and probed with a secondary horseradish peroxidase-conjugated antibody (Abcam) at a dilution of 1:15,000 in 1x PBST. After subsequent washing, the membrane was developed with an ECL SuperSignal West Pico kit (Pierce). After detection, the membrane was stripped and re-probed with mouse tubulin-alpha ab-2 (Sigma-Aldrich), 1:5000 dilution.

### Spontaneous Acrosome Reaction in the Presence of 10 μM DEHP in Capacitated Spermatozoa

Sperm was collected as described above. After 60 minutes of capacitation, spermatozoa suspension was transferred to clean micro tubes and centrifuged at 300 × *g* for 5 min at room temperature. The cells were then fixed in 4% PFA in 1X PBS for 15 minutes. At the end of fixation, an equal volume of 0.1 M ammonium acetate was added. The microfuges were centrifuged at 800 × *g* for 5 minutes. The supernatant was removed, and sperm cells were re-suspended in the remaining 100 μL. 30 μL of sperm suspension was spotted onto non-charged microscope slides and gently spread out with a glass pipette. The samples were allowed to air-dry for 15 min. Subsequently, the slides were washed in water followed by a methanol wash, and then water again, each wash step was done for 5 minutes. The slides were subsequently submerged in Coomassie brilliant blue (Sigma-Aldrich) solution for 2 minutes (0.11 g Coomassie brilliant blue, 20 mL water, 25 mL Methanol, and 5 mL glacial acetic acid). The slides were then rinsed in water to remove excess Coomassie and mounted with Mowiol mounting medium (Millipore).

### Flow Cytometry

Sperm aliquots (3 × 10^6^) were taken at 60min of capacitation and fixed in 3.7% PFA in PBS for 10 min at room temperature. After centrifugation at 500 × *g* for 10 min, spermatozoa were washed and subsequently permeabilized in 0.1% Triton X-100 for 10 min at RT. Non-specific binding sites were blocked by 0.1% BSA in PBST for 30 min at RT. Tyrosine phosphoproteins were localized using an anti-phosphotyrosine antibody, clone 4G10 (10 μg/ml) conjugated with CF 647 dye (Biotium) in PBS with 0.1% BSA for 1 hour at RT. Labeled spermatozoa were then washed in PBS and re-suspended in 250 μL PBS for flow cytometric analysis. 10,000 cells per sample were analyzed. Sperm fluorescence was quantified using the BD LSR Fortessa flow cytometer. FlowJo™ Software was used for data analysis.

### ROS production detection by Chemiluminescence assay

Spermatozoa were recovered by removing both caudae epididymides and placing them into a 30 mm Petri dish containing HS media. Spermatozoa were allowed to swim out for 15-20 min. Subsequently, the sperm suspension was equally divided between three Eppendorf tubes and spun down at 300 × *g* for 7 min. After the removal of the supernatant, the cells were re-suspended in an equal volume of HTF containing either ethanol or DEHP. The control tube contained 1 μL/mL ethanol and the treatment tube contained either 10 μM or a 100 μM DEHP. The suspensions were then capacitated at 37°C in 5% CO_2_ for 60 min. Detection of reactive oxygen species generated by sperm cells was done using the chemiluminescent agent - luminol following a previously described procedure(Agarwal et al. 2008). In brief, the chemiluminescent probe, luminol (Sigma-Aldrich, A8511-5G.) was freshly prepared before each experiment. After 60 min of capacitation, the samples were spun down at 300 × *g* for 7 min and re-suspended in 125 μM luminol in DPBS. Negative control, test sample, and positive control were prepared. 100 μL of 30% hydrogen peroxide solution was added to the positive control. A 100 μL aliquot of the cell suspension was taken from each sample for sperm count. The samples were then taken for Chemiluminescence measurements using the Lumicycle 32 (Actimetrics, Inc. Wilmette, IL). The luminometer measured Chemiluminescence at 37°C for 5 minutes. ROS production was expressed as counted photons per minute (CPM)/10^6^ sperm. Data were recorded using Actimetrics Lumicycle Data Collection software and analyzed using the Actimetrics Lumicycle Analysis program.

### Statistical analyses

For statistical analyses used in the manuscript the GraphPad Prism 5 software (GraphPad Software, Inc., La Jolla, CA) was used. Unpaired t-test was used to determine statistical significance for embryo survival, *IVF* and Chemiluminescence experiments, and assigning *p* ≤ *0.05* as the limit. Paired t-test was used for the AR, PY and flow cytometry experiments. All results are shown with the standard error of the mean. The significance of changes are indicated as follows: * *p* ≤ *0.05*, ** *p* ≤ *0.01*, *** *p* ≤ *0.001*

## Acknowledgments

This work was supported by March of Dimes, Pew Biomedical Scholars Award and Packer Wentz Endowment Will (to PVL), as well as Male Contraceptive Initiative grant to LGK and Summer Research Fellowship to JDR. We thank Dr. Andrew Modzelewski for his valuable advice and technical assistance with embryo work.

## Authors’ contributions

LGK and PVL conceived the project, designed the experiments and wrote the manuscript. LGK performed all studies, data acquisition and analysis for the manuscript. JDR helped with acrosome reaction, tyrosine phosphorylation, Chemiluminescence and flow cytometry studies. NP assisted with flow cytometry experiments and JM and LJK helped with Lumicycle and data analysis of luminol studies. All authors discussed the results and commented on the manuscript.

